# Large-scale expansions and replication stalling of Friedreich’s ataxia GAA repeats in an experimental mammalian system

**DOI:** 10.1101/2022.07.04.498737

**Authors:** Anastasia Rastokina, Jorge Cebrián, Nicholas Mandel, Rula Zain, Massimo Lopes, Sergei M. Mirkin

**Author notes:** A.R. and J.C. contributed equally to this work.

## Abstract

Human disease Friedreich’s ataxia (FRDA) is caused by large-scale expansions of (GAA)n repeats in the first intron of the *FXN* gene. While repeat expansions during intergenerational transmissions are causative for the disease development, somatic expansions additionally contribute to the disease progression. We and others have previously shown that (GAA)n repeats transiently pause the replication fork progression in cultured human cells. However, whether and by which mechanisms fork stalling underlies repeat expansions remained unclear. Here we developed a new genetically tractable experimental system to simultaneously analyze repeat-mediated fork stalling and large-scale repeat expansions in cultured human cells. It is based on a mammalian/yeast shuttle vector that can transiently replicate from the SV40 replication origin in human HEK-293T cells or be stably maintained in *S. cerevisiae* utilizing ARS4-CEN6; it also contains a cassette for selecting repeat expansions in yeast. Repeat expansions accumulate in mammalian cells and are then detected upon plasmid transformation into yeast. We found that large-scale expansions of (GAA)n repeats do occur in this experimental mammalian system. Further, we observed that repeat expansions’ frequency depends on several previously implicated proteins in replication fork stalling, reversal, and restart. These proteins include SHPRH, RAD52, ZRANB3, DDX11, SMARCAL1, HLTF, RECQ1 and WRN. Therefore, we propose that GAA repeat expansions might occur as a consequence of deregulated replication fork regression and restoration process.

## 2 Introduction

Expansions of simple, tandem DNA repeats lead to the development of ∼50 hereditary diseases in humans ^1–4^. One of these diseases is Friedreich’s ataxia (FRDA): a rare, autosomal recessive degenerative disease caused by the expansion of GAA repeats in the first intron of the frataxin gene (*FXN*) ^5^. FRDA patients have 70 to more than 1000 GAA repeat tracts, most commonly 600–900 on both alleles of the *FXN* gene ^6^. Lengthening of GAA repeat tracts results in a progressive reduction of the *FXN* mRNA ^7–9^, frataxin deficiency, mitochondrial dysfunction and cell death ^10, 11^. Consequently, the severity and the onset of the diseases are directly correlated with the length of the GAA repeat ^12^.

The mechanism of GAA expansions remains to be understood. Both *in vitro* and *in vivo*, (GAA)n repeats can adopt two unusual DNA structures: an intramolecular triplex called H-DNA ^13–17^ and a less characterized structure called sticky-DNA, which involves two distant (GAA)n repeats ^18, 19^. It was hypothesized that the formation of triplex DNA could be responsible for GAA repeat expansions ^20, 21^. An argument supporting this hypothesis comes from the genome analysis of an FRDA patient with a very mild and late-onset disease who has the (GAAGGA)64 repeat in the *FXN* gene, a repeat which cannot form H-DNA and is stably maintained in model experimental systems ^22^.

DNA triplexes formed by (GAA)n repeats block DNA polymerization *in vitro* ^13, 23^ and stall the replication fork progression in every experimental system studied, including bacteria ^24^, yeast ^25, 26^ and human cells ^27–29, 30^. We have previously developed a unique experimental system to detect and measure the rate of large-scale expansions of (GAA)n repeats in yeast ^31^. A subsequent unbiassed, genome-wide genetic screen identified several dozens of genes affecting the rate of repeat expansions, most of which encoded replication fork components ^32^. Replicative DNA polymerases and proteins involved in Okazaki fragment maturation strongly counteract repeat expansions ^33–35^. Finally, stabilization of H-DNA formed by the (GAA)n repeat by an RNA transcript (H-loop) additionally increases repeat instability ^36^. Altogether, these data led us to propose that GAA repeat expansions in yeast occur while the replication fork struggles to progress through the structure-prone DNA element.

This paper is aimed to study whether large-scale expansions of GAA repeats in human cells also occur during DNA replication. To this end, we created an experimental system that allowed us to analyze DNA replication of GAA repeat and their large-scale expansions in cultured human cells. It is based on a shuttle vector that utilizes extremely efficient T-antigen driven replication from the SV40 origin in human cells. It also contains our previously described cassette for selecting repeat expansions in yeast and can be stably maintained in *S. cerevisiae* ^31^. Repeat expansions accumulated during replication of this vector in mammalian cells are then detected upon its transformation into yeast. We found that GAA repeats cause stalling and reversal of the replication fork in human cells.

Furthermore, large-scale expansions of (GAA)n repeats efficiently occur in this system, making it the first experimental model to analyze them in human cells. We then conducted a candidate gene analysis of large-scale repeat expansions using the siRNA-mediated gene silencing approach. The depletion of 8 of the 19 proteins tested significantly impacted the frequency of GAA repeat expansions. Notably, those proteins were previously implicated in the unwinding of triplex DNA, fork reversal, and fork restoration. We hypothesize, therefore, that GAA repeat expansions in our system occur as a result of impaired replication fork progression through this hard-to-replicate DNA sequence.

## 3 Results

### Experimental system to study genome instability mediated by (GAA)n repeats in cultured mammalian cells

A new system that allowed us to measure large-scale expansions of (GAA)n repeats and repeat-mediated fork stalling in cultured human cells is presented in Figure 1A. It is a shuttle vector that encodes Tag for its efficient replication from the SV40 origin in human cells, an ARS4-CEN6 block for its stable maintenance in yeast, *S. cerevisiae,* and ColE1 replication origin to conduct cloning in *E. coli*. It also contains our previously described cassette to detect large-scale repeat expansions, which consists of an artificially split *URA3* reporter gene carrying the (GAA)100 repeat in its intron followed by the *TRP1* gene (*UR*-GAA100-*A3 TRP1* in Figure 1B). The addition of 10 or more repeats within the intron abrogates splicing of the *URA3* reporter, rendering yeast resistant to 5-fluoroorotic acid (5-FOA^r^). This vector was first transfected into human HEK-293T cells, where it was allowed to replicate for 48 h. Plasmid DNA was then isolated, treated with the *Dpn*I restriction enzyme to remove unreplicated DNA, and transformed into yeast, which serves as a read-out for repeat expansions that occur in mammalian cells. Note that only DNA fully replicated in human cells will withstand *Dpn*I treatment and, thus be able to transform to yeast. The frequency of repeat expansions was then estimated by comparing the number of 5-FOA^r^ clones with the total number of Trp^+^ transformants (Figure 1C). A similar methodology has previously been developed in the Lahue lab to analyze mid-scale expansions of the (CAG)25 repeat ^37–39^.

**Figure 1.**
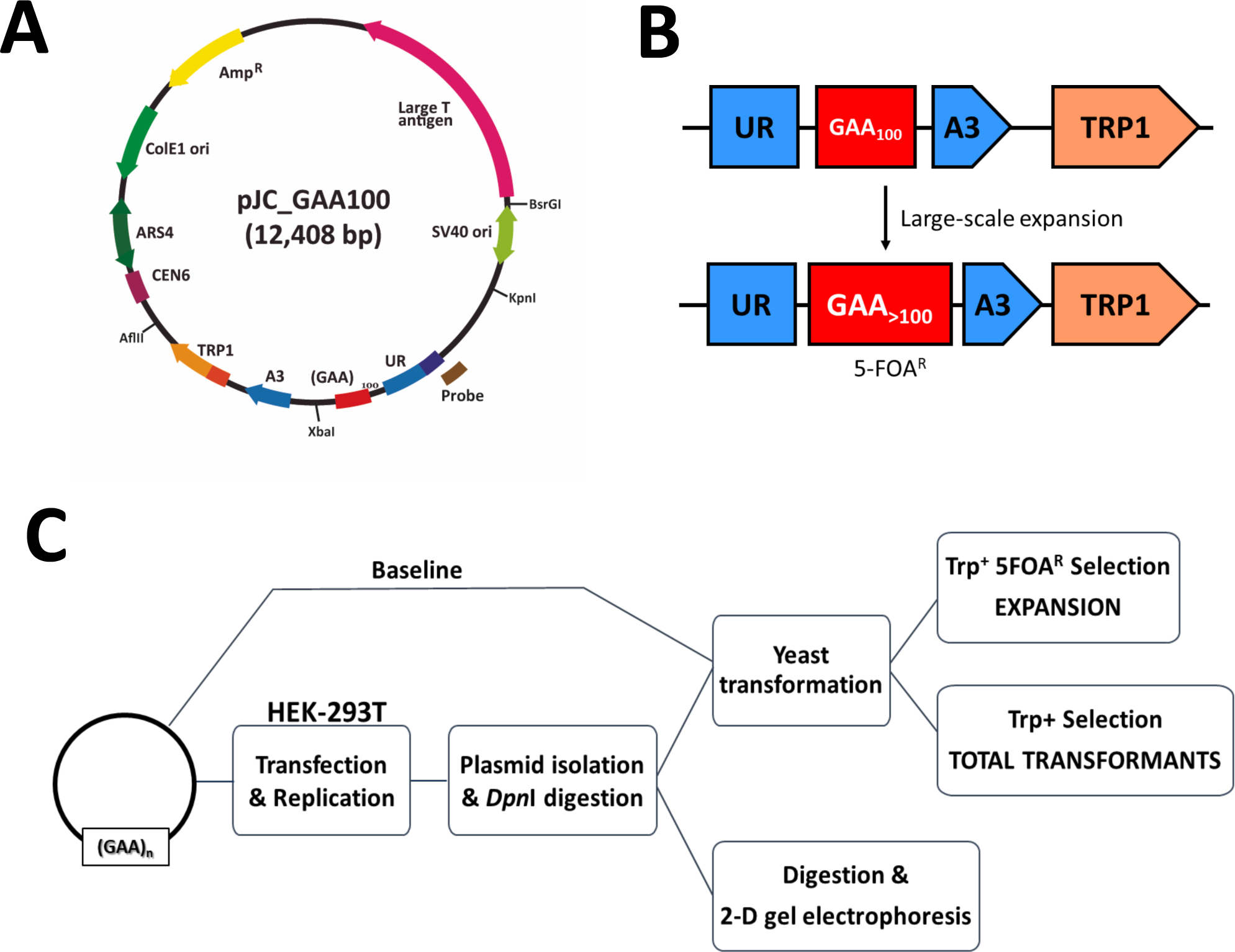
An experimental system to study genome instability and fork stalling caused by (GAA)100 repeats. Large-scale GAA-repeat expansions can be recovered from mammalian cells and analyzed in budding yeast. **(A)** pJC_GAA100 - Plasmid used in the assay. The relative positions of its most relevant features are indicated inside: The centromeric sequence CEN6, the Autonomous Replication Sequence (ARS4), the ColE1 unidirectional origin (ColE1 Ori), the ampicillin resistance gene (Amp^R^), the Large T antigen gene, the SV40 origin of replication (SV40 ori) and the selectable cassette for repeat expansions (*URA3-TRP1*: Depicted in B). Outside, the relative positions of sites recognized by specific restriction endonucleases are indicated. The plasmid was named according to the lagging strand template. **(B)** Schematic of the system to select for repeat expansion in yeast (previously described in Shishkin et al. 2009). An artificially split *URA3* gene contains 100 GAA repeats such that expansion events abrogate splicing and result in resistance to a 5 fluoroorotic acid (5-FOA) medium. The addition of 25-30 repeats increased the overall length of the intron beyond the splicing threshold. The selectable cassette was cloned into the pJC_GAA100 plasmid. *TRP1*: an auxotrophic marker for the selection of strains bearing the plasmids. **(C)** Schematic representation of the assay. Plasmids for the study were transfected into HEK-293T cells, and after culturing them for 48 h, DNA was isolated and digested with *Dpn*I. To study genetic instability, DNA was transformed into yeast. Colony PCR was performed to confirm expansion events. Expansion frequency was calculated by dividing the number of expansions by the total number of transformants. To calculate background expansion frequency, plasmids skipping mammalian cells transfection were transformed into yeast. To study the replication through the repeats, DNA was digested with restriction enzymes, and replication intermediates were analyzed by Two-Dimensional (2D) agarose gel electrophoresis.

We placed the homopurine run on the lagging strand template for human DNA replication (Figure 1A) because DNA triplexes formed by homopurine-homopyrimidine repeats strongly block replication when their homopurine runs are in the lagging strand template, making homopurine repeats more unstable in this orientation ^24, 25, 31, 40–45^.

Plasmids with SV40 replication origin and large Tag replicate very efficiently ^37, 46^ in human cells in the presence or absence of (GAA)n repeats ^27^. By and large, replication of these plasmids involves proteins of the mammalian replication fork, but there are two noticeable exceptions. First, Tag serves as a replicative helicase instead of the CMG helicase ^47–49^. Second, DNA polymerase δ (Pol δ) conducts both the leading and lagging strand DNA synthesis in those plasmids ^50^, whereas the syntheses of the leading and lagging strand during eukaryotic genome replication are carried out by DNA Pol ɛ and Pol δ, respectively ^47–49, 51^. As a result, SV40-driven replication is generally more prone to fork collapse and restart ^50^. While we are aware that these differences between the SV40-driven and regular DNA replication machinery can complicate the comparison of our data with the results from patient cell studies, we have chosen this experimental set up because the robustness of this replication system provided us with enough replication products to simultaneously analyze replication fork progression through the repeat as well as the instability of the repeat.

### Genetic analysis of GAA repeat instability

Plasmid DNA replicated in mammalian cells was treated with *Dpn*I and transformed into yeast. To determine the frequency and the scale of GAA repeat expansions, 5-FOA^r^ colonies were tested by single colony PCR (Figure 2A). The expansion frequency was calculated by dividing the confirmed expansions in the 5-FOA resistance plates by the total number of Trp^+^ transformants (Figure 1C) using FluCalc calculator ^52^. We compared these data with the baseline frequency of expansions that either originated in bacteria or in the process of yeast transformation by transforming bacterial plasmid DNA that was not replicated in human cells. Figure 2B shows the frequencies of repeat expansions in human embryonic kidney cells with or without the Tag gene (HEK-293T and HEK-293, respectively). Note that replication of HEK-293 cells is driven by the Tag encoded by our plasmid. One can see that the frequency of GAA repeat expansions in both cases significantly exceeds their baseline frequency.

**Figure 2.**
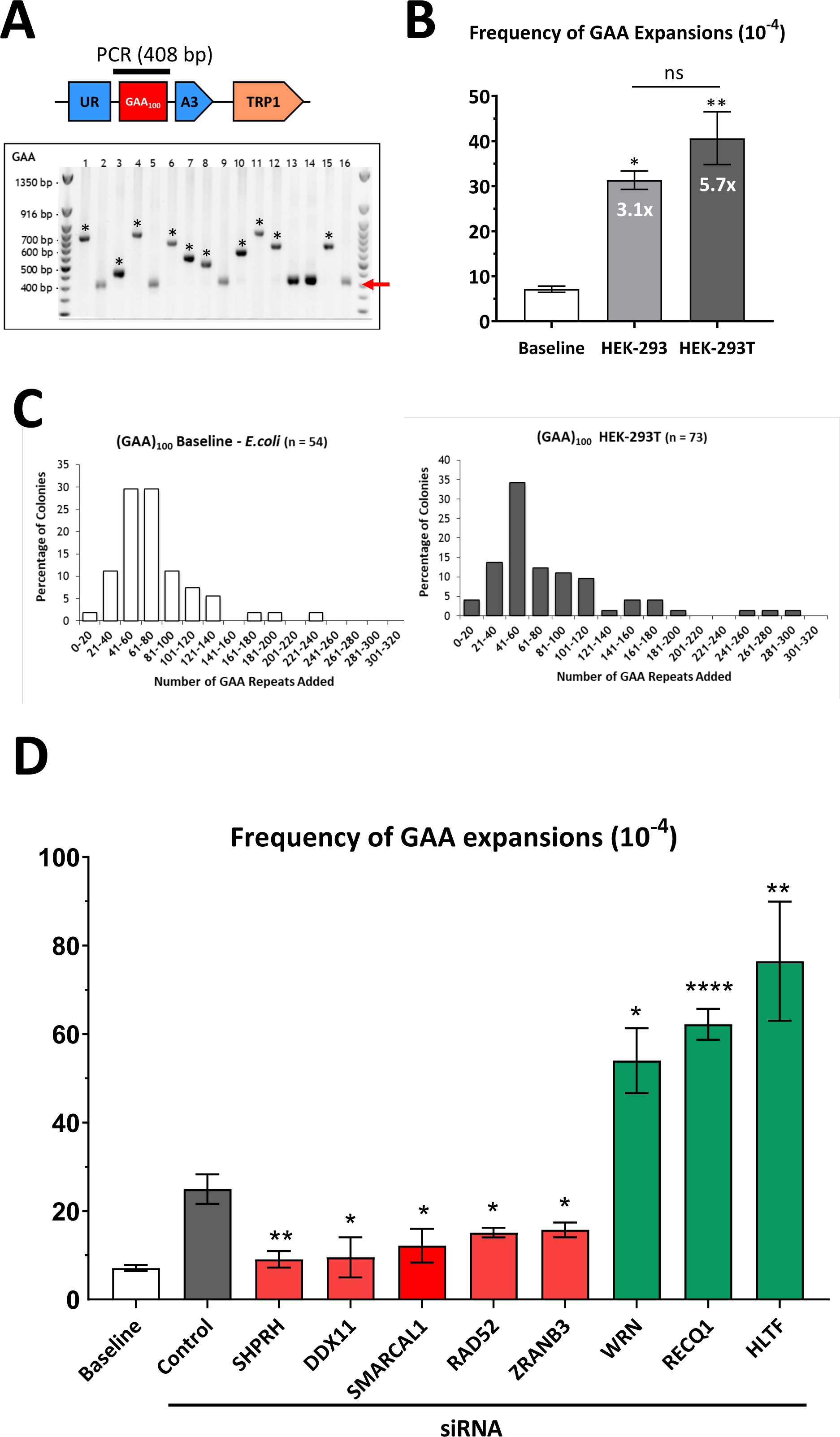
Frequencies of GAA repeat expansion after their replication in HEK-293T cells and upon candidate gene knockdown by siRNA. (A) PCR confirmation of GAA expansions in HEK-293T cells. 5-FOA-resistant colonies were analyzed from independent transformations. On the upper part, a cartoon representation of a cassette with a bar above the GAA repeats indicates the product of single-colony PCR, used to determine the repeat length. The red arrow points to the initial length of (GAA)100 repeats. Only clones with expansions were used to calculate expansion frequencies in Figure 2D. ‘*’ in lanes denote large-scale GAA expansions. **(B)** Expansion frequencies for pJC_GAA100 plasmid derived from *E. coli* (baseline), HEK-293, and HEK-293T cells. Error bars indicate the standard error of the mean. ‘*’ P<0.05, ‘**’ indicates a P<0.001 two-way Welch ANOVA test **(C)** Distribution of repeats added to (GAA)100 for plasmids HEK-293T cells (dark grey bars). The geometric mean of repeats added is 65 repeats (interquartile range 50.5-84.5) for the *E. coli*- baseline and 65 repeats (interquartile range 44.5-100.5) for HEK-293T. **(D)** The baseline expansion frequency upon yeast transformation by plasmid DNA isolated from *E. coli* is shown (white bar) for the comparison. Error bars indicate the standard error of the mean. Significance compared to the siControl frequency value was determined using a two-way Welch ANOVA test, ‘#’ P<0.05 versus baseline, ‘*’ P<0.05, ‘**’ indicates a P<0.001 versus siControl. See Table S1 for details.

The expansion frequency was slightly higher when plasmids replicated in HEK-293T cells than in HEK-293 cells (Figure 2B), which is likely due to instant plasmid replication from Tag expression in HEK-293T cells, which occurred prior to transfection. This difference, however, was not statistically significant. Notably, the *hMLH1* gene in the HEK-293T cells is epigenetically silenced by promoter hypermethylation, while it is functional in the HEK-293 cells ^53^. Since there was no significant difference in GAA expansion frequency between the two cell lines, we can rule out that MutL complex plays a role in this process (Figure 2B). This observation contrasts earlier data from a transgenic mice model stipulating the role of MutL complex in triple repeat expansions during intergenerational transmissions ^54^. We attribute this difference to the fact that repeat expansions in the transgenic mice model are small scaled in nature (just a few repeats are added per intergenerational transmission).

Unlike previous models, in our system the mean number of added repeats corresponds to 65 repeats (Figure 2C), and we were able to detect up to 300 added repeats when the plasmid replicated in HEK-293T cells. To the best of our knowledge, this is a unique system to observe large-scale expansions after only 48 h in human culture cells.

To identify proteins involved in large-scale GAA expansions, we used candidate-gene analysis. We and others have previously identified multiple genes and proteins involved in repeat expansions in various model systems ranging from yeast ^25, 26^ to human cells ^27–29^. Evidently, different proteins are involved depending on the repeat type, expansion scale, cell type and organism (reviewed in ^1, 55–57^). Notably, most of those proteins are the components of DNA replication and repair. As discussed above, it does not seem that mismatch repair significantly affects GAA repeat expansions in our system (Figure 2B). Therefore, we selected 18 candidate genes that encode for proteins involved in DNA replication and post-replication repair, some of which were also implicated in GAA repeat instability in yeast. Those genes were knocked down using pooled siRNAs (Figure S2A, B), followed by measuring repeat expansion frequency.

Our candidate genes can be divided into several functional groups. The first group of genes – FEN1, TIMELESS, and CLASPIN – encode the replication fork components previously implicated in repeat expansions and instability. Flap endonuclease 1 (Fen1), which is required for the flap-removal during Okazaki fragments maturation and is involved in various DNA repair pathways, was shown to prevent expansions of multiple repeats in a yeast experimental system ^1^, while its role in mammalian cells remains questionable ^58, 59^. TIMELESS (yTof1) and CLASPIN (yMrc1) proteins, which are components of the fork–stabilizing complex, were shown to prevent CAG repeat instability and expansions in yeast and human cells ^60, 61^, as well as GAA repeat instability in yeast ^31, 32, 61^. In our system, however, siRNA knockdown of these proteins did not affect GAA expansion frequency (Figure S3).

The second group of genes – ATR (yMEC1) and ATM (yTEL1) – trigger DNA damage response (DDR) caused by the replication stress. They were previously shown to stabilize CAG•CTG repeats in yeast, mice and human cells ^62^. siRNA knockdown of those genes in our system did not show a statistically significant effect on GAA repeat expansions, albeit the depletion of ATM slightly (1.6-fold) elevated repeat expansions (Figure S3).

The third group of genes –RAD51, RAD52, BRCA1, and BRCA2 – encodes the key components of homologous recombination (HR) and DSB-repair in mammals ^63, 64^. Homologous recombination has been implicated in both the promotion and suppression of repeat expansions and instability in various experimental systems ^65^. In our case, knocking down RAD51, BRCA1 or BRCA2 did not affect GAA repeat expansions. At the same time, knocking down RAD52 significantly decreased GAA repeat expansions (Figure 2D). Besides canonical HR ^66–68^, RAD52 was also implicated in the pathway of DSB repair break-induced replication (BIR) ^69, 70^. BIR, and specifically RAD52, were shown to promote CAG and CGG repeat expansions in yeast and cultured human cells ^71, 72^. POLD3 (yPol32), a small subunit of DNA polymerase δ, is an essential protein required for DNA synthesis during BIR ^73^. In our system, the depletion of POLD3 did not change the frequency of GAA repeat expansions, effectively ruling out the role of BIR. Finally, in some studies, RAD52 protects reversed replication forks from degradation by exonucleases ^74^. We suggest that this function of RAD52 is essential for GAA repeat expansions in our case (see below and Discussion).

The fourth group of genes – HLTF and SHPRH – encode ubiquitin ligases involved in the poly-ubiquitination of PCNA critical for DNA damage tolerance, specifically fork reversal and template switching ^75, 76^. The yeast homolog of these genes, RAD5, was shown to promote GAA and ATTCT repeat expansions ^31, 77^. In addition, the helicase activity of HLTF has been implicated in fork reversal ^78^. The knockdown of HLTF, but not SHPRH, has been shown to elevate CAG repeat expansions several folds in human cells ^79^. Here we show that HLTF knockdown increases GAA repeat expansion frequency (Figure 2D), similarly to what was observed for CAG repeats. Surprisingly, the knockdown of SHPRH dramatically reduced GAA repeat expansion frequency (Figure 2D).

The fifth group of genes –SMARCAL1 and ZRANB3 –encode SWI/SNF helicases and ATPases, which catalyze replication fork reversal and restart ^80–84^. Consequently, depletion or inactivation of these proteins hinders the ability of a replication fork to recover from replication stress, particularly by increasing the frequency of double-strand breaks ^80, 85–87^. We were interested in these proteins because of their role in replication fork reversal and restart, the role of which in promoting repeat expansions in various experimental systems has been widely discussed ^1, 27, 88, 89^. In line with those data, depletion of either SMARCAL1 or ZRANB3 proteins prevented GAA repeat expansion in our system (Figure 2D).

The sixth group, DDX11 and FANCJ, are members of the DEAD/DEAH box helicase family ^90, 91^ that prevent replication stress and mediate HR repair by directly interacting with Pol δ ^92, 93^. We were specifically interested in the DDX11 helicase, as it has been shown to unwind triplexes in vitro ^94, 95^. Here, we show that the depletion of DDX11 by siRNA dramatically decreased repeat expansions (Figure 2D). At the same time, depletion of FANCJ did not cause any change in GAA expansion frequency (Figure S3).

The next group is the RecQ family of DNA helicases that consists of five homologs: RECQ1, WRN, BLM, RECQL4, and RECQL5, which has been shown to be is a significant family in fork restart. RecQ helicases interact physically and functionally with PCNA, RPA and DNA polymerase δ ^96–101^. RECQ1 itself has a 3′-5′ directed DNA unwinding capacity that helps maintain genomic integrity by preferentially restoring reversed forks to their original three-armed configuration *in vitro* and *in vivo* ^102^. Here, we show that the depletion of RECQ1 increases the (GAA)n repeat expansion in our system. One of the most well studied members of the RecQ family, Werner (WRN), is involved in resolving a variety of DNA substrates: replication forks, flaps, D-loops, bubbles, Holliday junctions, and G-quadruplexes (G4) ^103, 104^. The depletion of WRN increases repeat expansion in our system, similarly to the depletion of RECQ1 and HLTF.

Besides the RecQ family of helicases, DNA2 also is involved in reversed fork processing. Human RECQ1 has been shown to limit DNA2 activity by preventing extensive nascent strand degradation at the reversed fork. Previously, it has been shown that DNA2 mediates DNA end resection together with WRN. However, the depletion of DNA2 did not influence repeat expansions in our system ^105^.

In summary, five proteins, SHPRH, DDX11, SMARCAL1, RAD52 and ZRANB3, appear to promote GAA repeat expansions in our experimental human cell system, while three proteins, HLTF, WRN and RECQ1, appear to counteract them (Figure 2D, S3, and Table S1).

### Replication fork progression through the (GAA)100 repeat

We and others have previously shown that expanded (GAA)n repeats stall replication fork progression in yeast and human cells ^25, 27–29^. Notably, however, quantification of the fork stalling in the SV40-driven mammalian episome, similar to that described here, has not been carefully analyzed. To fill in this gap, we measured the strength of the repeat-mediated fork stalling in our episome using 2-D electrophoretic analysis of replication intermediates. In brief, DNA plasmids with and without (GAA)n repeats were transfected into HEK-293T cells, replication intermediates were isolated 48 h post-transfection, treated with *Dpn*I to get rid of unreplicated DNA, digested by *Bsr*GI and *Xba*I restriction endonucleases and separated by 2-D agarose gel electrophoresis followed by Southern hybridization. This digestion deliberately positions the repeat-mediated stall site on the descending part of the replicative Y-arc (Figure 3B). Previously, we and others analyzed GAA-mediated stall sites positioned on the ascending portion of the Y- arc. Note however, that this setting results in a partial overlap of the stall site with the reversed fork (Figure S4), complicating the analysis of fork reversal. By placing the repeat on the descending half of the Y-arc, we were able to address this problem (see below).

**Figure 3.**
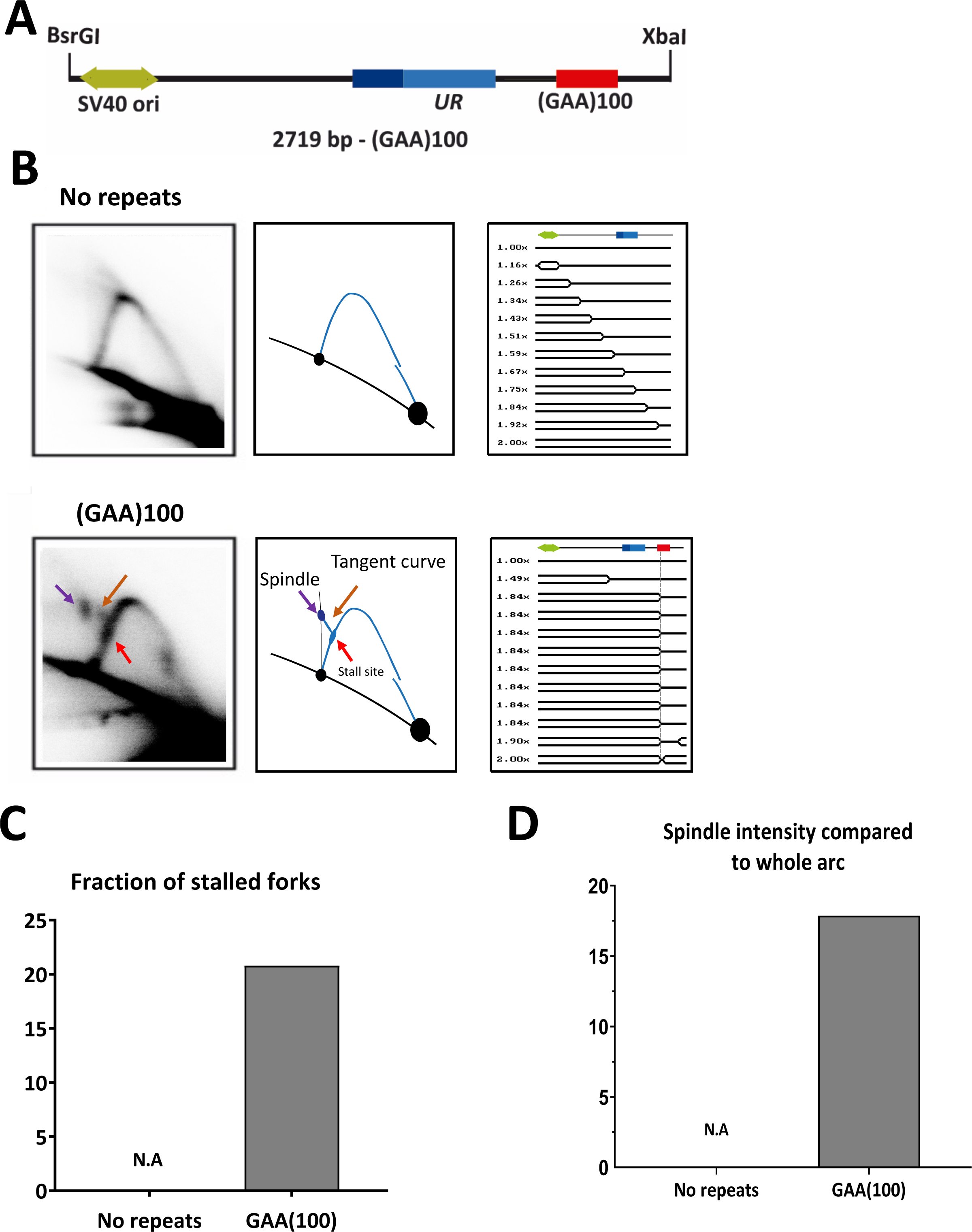
Analysis of pJC_GAA replication in HEK-293T cells by 2D gel electrophoresis. (A) Linear map of pJC_GAA100 restriction fragment. **(B)** Representative 2D gels of replication through (GAA)0 and (GAA)100 repeats are shown in the far-left column with their corresponding interpretative diagrams to their right. DNA was isolated, digested with *Bsr*GI, *Xba*I, and *Dpn*I, and analyzed by 2D gels. The simulation program 2D gel ^106^ was used to predict the shape of twelve consecutive replication intermediates (RIs). A linear map is shown on top of each series of RIs, showing the relative positions of SV40 ori (green) UR (blue). The red arrow points to the location of the stall due to the GAA 100 repeats **(C)** Quantification of the fraction of stalled forks. The ratio of radioactivity in the peak area to that corresponding area of a smooth replication arc reflects the extent of replication slowing. Quantification was done with Image J (NIH). Error bars indicate the standard error of the mean. N.A. = Non-Applicable. **(D)** Quantification of spindle spot intensity compared to the whole arc. The ratio of radioactivity signal at the spindle spot was compared to the radioactivity signal of the whole arc on Image Lab®. Error bars indicate the standard error of the mean. N.A. = Non-Applicable.

The simulation program for 2-D gels ^106^ was used to predict the shape of the Replication Intermediates (RIs) responsible for the patterns observed. The pattern detected for the control plasmid corresponded to that expected for unconstrained replication of the circular plasmid where the initiation occurs at SV40 origin in a bi-directional manner (simple-Ys’s shape) (Figure 3B). However, the repeat-containing plasmid produced a different pattern. In this case, replication would initiate at the SV40 origin bidirectionally, creating a bubble. The leftward moving fork progresses unconstrained, while the rightward moving fork stalls at GAA repeats, resulting in the accumulation of simple Y intermediates with a mass of ∼1.65X. In the event of complete stalling, replication must be completed by the leftward replication fork, leading to the accumulation of double-Y intermediates migrating above the descending arc. Alternatively, the rightward fork stalled at the repeat can regress or reverse, forming chicken foot intermediates, which should migrate somewhere in between the stall site and double-Y intermediates. The reversed fork can be subsequently restored, or the replication will be completed by the opposite replication fork.

Experimental data shown in Figure. 3B clearly demonstrates the presence of the stall site at the expected position on the descending arc, the tangent curve exiting from this stall and the spindle-shape spot corresponding to the array of double-Y intermediates with and without the chicken foot structure. These results confirm that GAA repeats strongly stall replication in HEK-293T cells in agreement with previous data ^27^. To quantify these results, we first normalized the signal at the stall site to the signal replication arc underneath (Figure S5). Quantification showed that ∼20% of all replication forks stall at the (GAA)100 repeats, while no stalling at this position was observed for the no-repeat plasmid (Figure 3C). The spindle-shaped spot was quantified by comparing its intensity to the rest of the Y-arc (Figure S1B). It was not present for the no-repeat plasmid but accounted for a ∼17% of replicative intermediates in the repeat-containing plasmid (Figure 3D).

We then analyzed whether proteins that affected GAA repeat expansion in our system (Figure 2D) affected replication fork progression through this repeat as well. To this end, we used 2-D electrophoretic analysis of RIs isolated from cells depleted of RAD52, ZRANB3, SMARCAL1, SHPRH, HLTF, DDX11, WRN and RECQ1 proteins via siRNA (Figure 4A). Note that depletion of several of these proteins changed the shape of the stall site on the descending arc: instead of the well-defined spot in the non-treated cells it converted to an elongated, rectangle-like spot (see ZRANB3, WRN, RECQ1, SMARCAL1 in Figure 4A). This made the comparison of the stall sites between different siRNA treatments ambiguous. Thus, we decided to analyze spindle-shaped intermediates that were present in every case. Quantification of those intermediates is shown in (Figure 4B and Table S2). Spindle-shaped spot intensity was strongly decreased upon SHPRH and ZRANB3 depletion, but increased upon DDX11, SMARCAL1 and RECQ1 depletion. Note that in the cases of ZRANB3 and SHPRH depletion, the spindle spot looks minimal: this signal pattern of replication intermediates exiting from the stall site as a tangent line corresponds to a replication fork that tries to reverse but fails.

**Figure 4.**
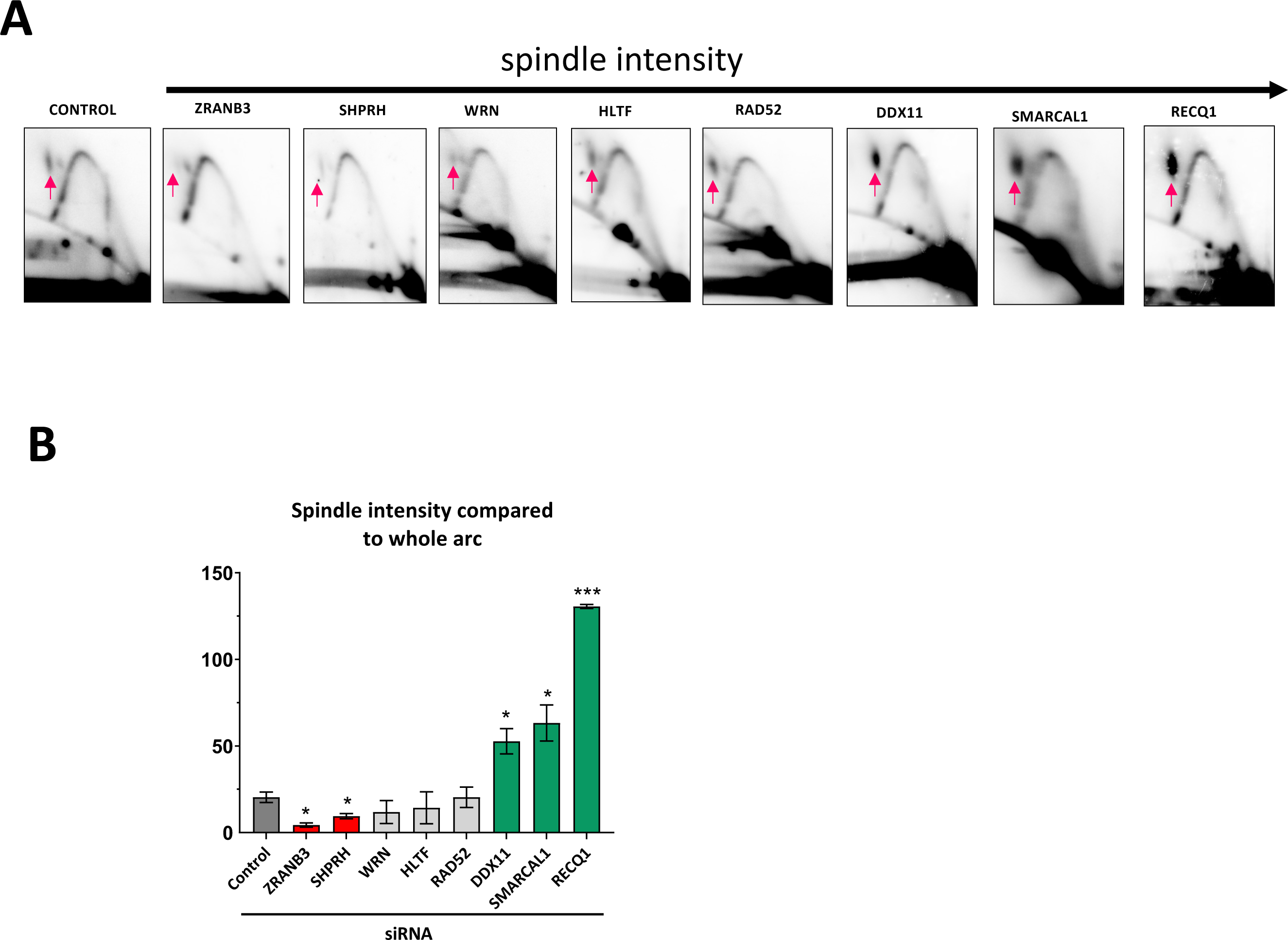
Analysis of replication through (GAA)100 repeats by two-dimensional (2D) agarose gel electrophoresis in HEK-293T cells. (A) Representative 2D gels of replication through (GAA)100 repeats in HEK-293T cells. DNA was isolated, digested with *BsrG1, DpnI* and *XbaI*, and analyzed by 2D gel. The red arrow points to the location of the stalling at the GAA repeats. At least three experiments were analyzed for each siRNA treatment. **(B)** Quantification of the fraction of stalled forks. The ratio of radioactivity in the peak area to that corresponding area of a smooth replication arc reflects the extent of replication slowing. Quantification was done with Image J (NIH). Error bars indicate the standard error of the mean. *‘\**’ indicates a *P*<0.05 two-way Welch ANOVA test. See Table S2 for details. (**C)** Quantification of the spindle spot at the replication fork. The ratio of radioactivity signal at the spindle spot was compared to the radioactivity signal of the whole arc on Image Lab®. Error bars indicate the standard error of the mean. ‘*’ *P*<0.05, ‘***’ indicates a *P*<0.0001 versus siControl. See Table S2 for details.

In sum, five out of the eight proteins that affected repeat expansions in human cells also changed the character of the replication fork progression through the GAA repeat. These data point to a link between replication and large-scale expansions of GAA repeats in human cells.

## 4 Discussion

In this study, we describe an experimental system to study GAA repeat instability in which repeat expansions that occurred during SV40-mediated episomal replication in human cells were detected upon transformation into yeast (Figure 1). Remarkably, we observed that large-scale repeat expansions did efficiently occur in this system: up to 300 repeats were added, while the mean number of added repeats was 65 (Figure 2C). While our experimental setting is similar to that previously developed in the Lahue lab for studying CAG repeat expansions ^37–39^, the latter experimental system was used to study mid-scale (up to 15 repeats) expansions. The only known instance where large-scale repeat expansions were observed in a mammalian experimental system was a specific transgenic DM1 mouse model ^107^, but the reasons for the big jumps of CTG repeats in these mice were unclear and never investigated further. Thus, our experimental system is unique for studying large-scale expansions in human cell lines.

We and others previously observed that GAA repeats cause replication fork stalling in various experimental systems, including SV40-based episomal replication ^25, 27–29^. In accord with these observations, we observed clear-cut replication fork stalling and reversal in our episome (Figure 3). This data prompted us to investigate various candidate genes that affected DNA replication and post-replicative repair and were previously implicated in repeat expansions in different experimental systems. Altogether, we analyzed nineteen candidates and found that siRNA depletion of eight candidates significantly affected the frequency of repeat expansions (Figure 2D). Strikingly, seven of those candidate genes (SHPRH, RAD52, RECQ1,SMARCAL1 WRN, HLTF and ZRANB3) were previously implicated in replication fork reversal/restart ^81–84, 102–104, 108–117^, while DDX11 is a DNA-helicase capable of unwinding DNA triplexes *in vitro*^94, 95^.

Using electrophoretic analysis of the replication intermediates, we then looked at changes in the replication fork progression through the GAA repeat upon depletion of the proteins encoded by these eight candidate genes. Specifically, we concentrated on the spindle-shaped spots that correspond to reversed forks intermediates and the double-Y intermediates formed by the opposite replication fork. Inactivation of SHPRH and ZRANB3 decreased, while DDX11, SMARCAL1 and RECQ1 depletion increased the intensity of these spindle-shaped spots.

Overall, our data points to the role of the replication fork stalling, reversal, and restart in GAA repeat expansions during episomal replication in human cells.

While fine molecular details of how fork stalling, reversal, and restart at the GAA repeat leads to its expansions are far from being clear, we can propose a tentative working model (Figure 5). During SV40 episomal replication, the T-antigen serves as a helicase, both the leading and lagging strands are synthesized by DNA Pol δ ^47, 118^, and there is a relatively poor coupling between the helicase (T-antigen) and DNA-polymerase delta (Pol δ) ^119^, making it prone to fork reversal and restart ^119, 120^. We suggest that a triplex transiently formed by the GAA repeat during DNA replication causes fork stalling (Figure 5A). For an already wobbly SV40 replication fork, this triplex can cause potent fork reversal. Fork reversal requires several redundant activities, including HLTF and SHPRH involved in the PCNA poly-ubiquitination and helicases ZRANB3 and HLTF. Typically, fork reversal also requires the activity of RAD51 protein ^121, 122^. Since we don’t see any effect from RAD51 depletion on repeat expansion, we may be dealing with a fork regression limited to the repetitive DNA, rather than a full-scale fork reversal. Depleting SHPRH and ZRANB3 reduces repeat expansions and decreases the spindle-like replication intermediates, implicating these proteins in the fork regression in our system (Figure 5B). Since RAD52 promotes repeat expansions in our system, we believe that it stabilized the regressed fork by preventing its excessive reversal and degradation ^74^.

**Figure 5.**
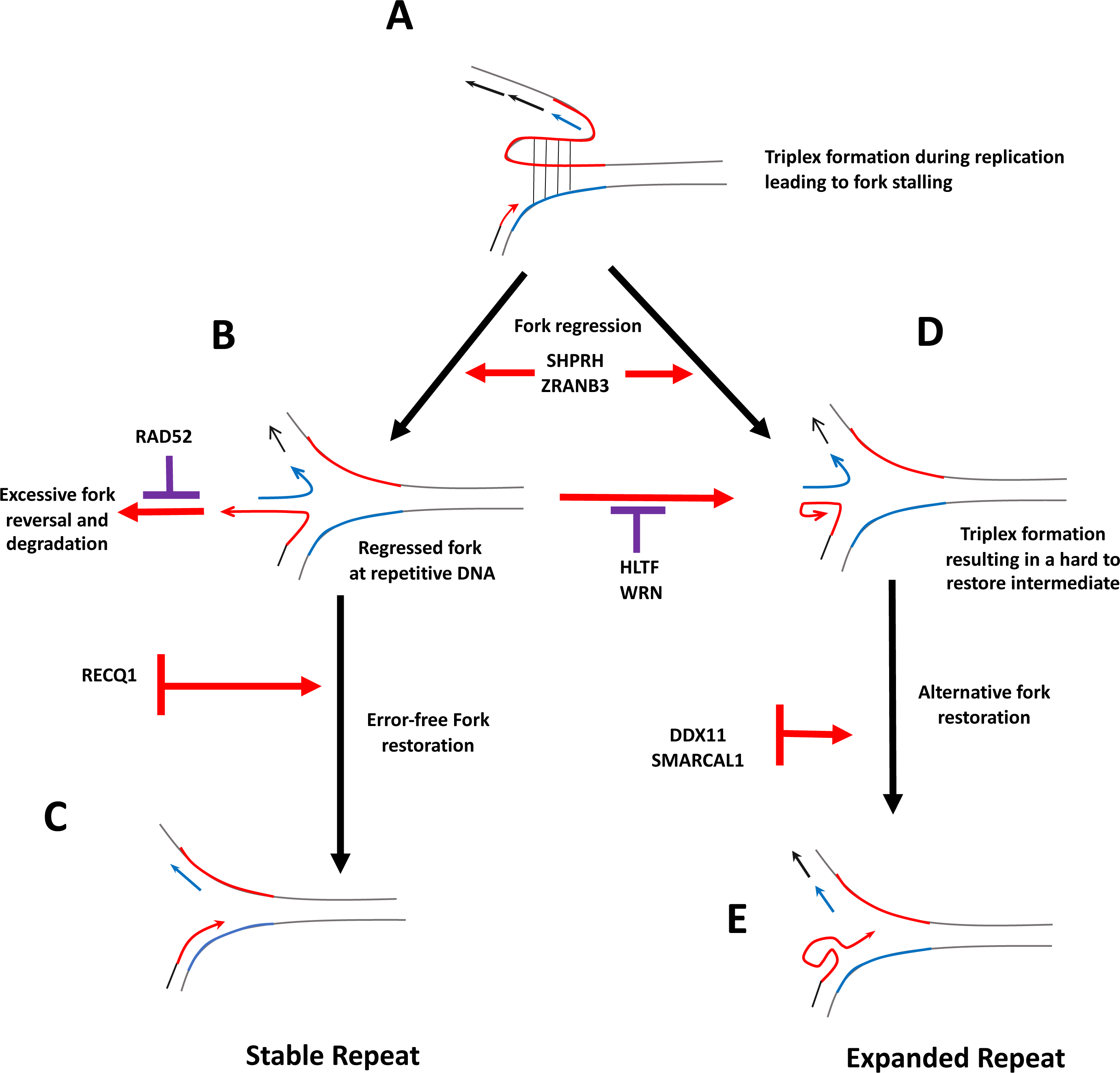
Proposed model for GAA large-scale expansions. (A) (GAA)100 repeat (in red) form a triplex during the replication halting replication **(B)** Fork back up is initiated by SHPRH and ZRANB3. This fork backup can be degraded if RAD52 is not present to protect the fork. **(C)** Subsequently, the fork backup is restored via an error-free fork mechanism with the help of RECQ1 and WRN. **(D)** Fork back up could alternatively fold back, forming a triplex with the nascent leading strand making this structure hard to retore intermediate. HLTF helicase, by binding to the 3’overhang via its HIRAN domain, could counteract this triplex. **(E)** This triplex formation could be resolved by an alternative, less precise pathway of fork restoration with the use of DDX11 helicase, causing repeat expansion due to an out of register realignment of repetitive DNA.

The fork regressed at the repeat is subsequently restored by the RECQ1 helicase ^84, 102, 123^, resuming normal replication without adding extra repeats (Figure 5C). Consistent with this idea, the depletion of RECQ1 leads to the accumulation of spindle-shaped intermediates (Figure. 5B) and increased repeat expansions. Alternatively, a repetitive single-stranded portion of the nascent leading strand can fold back, forming a triplex (Figure 5D). HLTF and WRN helicases are expected to counteract this process, since they are known to bind to single-stranded 3’-overhangs ^111, 115, 116, 124–127^. Indeed, depletion of both helicases increased repeat expansions in our system. If, however, this triplex is not unraveled by the concerted action of the above helicases, an alternative and less precise pathway of fork restoration, which involves DDX11 and SMARCAL1, might take place. Both DDX11 and SMARCAL1 are helicases that are known to efficiently unwind various non-B DNA secondary structures, including triplex DNA ^80, 84, 92, 93^. We propose that during this alternative pathway of fork restoration, complementary strands of the repeat can realign out-of-register, ultimately leading to an expansion (Figure 5D). This idea is consistent with our observations that depletion of DDX11 and SMARCAL1 decreases repeat expansions and leads to a significant accumulation of spindle-shaped intermediates, possibly corresponding to structures shown in Figure 5D.

How do our data and model compare to other studies of GAA repeat expansions? The discussion below concentrates on the data from the genetically tractable systems rather than from human patient data.

Baker’s yeast is the only other system where large-scale expansions of GAA repeats were studied experimentally ^26, 31, 52, 72, 77, 128^. It was firmly established that in dividing yeast, large-scale expansions of GAA repeats (∼60 repeats at a step) occur during DNA replication ^31, 32, 128^. They likely happen during Okazaki fragment maturation in the course of the lagging strand synthesis ^34^. R-loop formation additionally promotes expansions at the GAA repeat ^32, 36^ or checkpoint activation ^129^. The only study of mid-range GAA repeat expansions (10-to-15 repeats per generation) in yeast showed that they are promoted by malfunctioning of the CMG helicase followed by BIR ^130^.

As for mammalian system, Ed Grabczyk observed progressive accumulation of GAA repeats in a HEK-293T cell line depending on the activity of MutL gamma ^131^. Note, however, that only 1-to-2 repeats were added per cell generation in his system. In a transgenic FRDA mice line, small-scale intergenerational expansions of GAA repeats were inhibited in the presence of a mismatch repair system ^54^, while somatic expansions were promoted by the MMR ^132^.

The conclusions of our study differ from those in sensitive details: (i) MMR (or at least the MutL complex) does not seem to play a role; (ii) our data argue against expansions during the lagging strand DNA synthesis; and (iii) BIR does not seem to be involved.

At the same time, our data/interpretations are consistent with the study of Gerhardt and co-authors ^28^ analyzing replication of GAA repeats in FRDA patients’ cells. They found that expanded GAA repeats in iPSC derived from FRDA patients cause profound replication fork stalling. Further, preventing triplex formation by GAA repeats via specifically designed polyamides decreased fork stalling and increased repeat stability.

Also, there are evident similarities between our data and the data on the mid-scale CAG repeat expansions in yeast and mammalian cells that implicated fork reversal and restart in the process ^60, 79, 133–136^. It is foreseeable that in our case, starting expansions of GAA repeats that occur during fork reversal and restart, become very large during subsequent rounds of episomal replication.

In summary, we developed a first of a kind, genetically tractable experimental system to study large-scale expansions of FRDA GAA repeats in cultured human cells. Our candidate gene analysis implicates fork regression and restoration in this process. We believe that this system could be a valuable tool to evaluate the efficiency of perspective FRDA drugs targeting the instability of GAA repeats.

## 5 Methods

### Plasmids

The plasmid pJC_GAA100 was constructed by conventional cloning methods in several steps using the pLM113 plasmid as a backbone [1]. First, pRS316 was digested by *Sal*I and *Sac*I to obtain the ARS4-CEN6 sequence, and it was inserted between *Sal*I and *Sac*I of pML113 (gift from M. Lopes), creating a centromeric plasmid called pJW12 (8835 bp). The pJC_GAA100 plasmid (12408 bp) was obtained by inserting the repeat-containing-URA3 cassette *Ale*I-*Stu*I fragment of pYes3-T269-GAA100 [2] into the blunt-ended *Eco*RV site of pJW12. Note, in our pJC_GAA100 plasmid, GAA repeats comprise the template for lagging strand synthesis regarding the SV40 origin. The pJC_GAA0 (No repeats) plasmid was obtained with the same approach as GAA100, but the inserting *Ale*I-*Stu*I fragment was from pYES-TET644 [2]. All plasmid constructs were isolated from the *E.coli* SURE strain (Stratagene) and (GAA)100 repeats were confirmed by Sanger sequencing.

### Cell culture, transfection, and siRNA treatment

HEK-293T (ATCC) were grown in Dulbecco’s modified Eagle medium (DMEM, Gibco) supplemented with 10% fetal bovine serum (FBS) and MycoZap^TM^ Plus-CL (Lonza). Cells were transfected with pJC_GAA100 plasmid by using JetPRIME^®^ (Polyplus-transfection) according to the manufacturer’s instructions. Briefly, cells were seeded on day 0 and transfected on day 1 with siRNA. On day 2, cells were transfected with the GAA100 shuttle vector and retransfected with siRNA. After another two days, DNA was isolated to measure expansion frequencies or to perform Two-Dimensional (2-D) agarose gel electrophoresis. Gene silencing was confirmed by Western blotting. Figure S2 shows the details regarding siRNAs and antibodies used in this study.

### DNA isolation

Plasmid DNA was recovered 48 h post-transfection by a modified Qiagen Miniprep protocol as described in [1]. Briefly, cells were washed with PBS and then resuspended in Qiagen Buffer P1 and lysed in 0.66% sodium dodecyl sulfate, 33 mM Tris-HCl, 6 mM EDTA, 66 μg/ml RNase followed by digestion with 0.5 mg/ml proteinase K for 90 min at 37°C. Samples were subject to brief, 30 s, base extraction with 0.75 ml 0.1 M NaOH and proteins precipitated by addition of Qiagen Buffer P3 (4.2 M Gu-HCl, 0.9 M potassium acetate pH 4.8). Cell debris was pelleted at 29,000*g* for 45 min and supernatant loaded onto a Qiagen Miniprep spin column. Columns were washed with Qiagen Buffer PB (5 M Gu-HCl, 30% ethanol, adding 10 mM Tris-HCl, pH 6.6) and 0.75 ml Qiagen Buffer PE (10 mM Tris-HCl, pH 7.5, 80% ethanol) and plasmid DNA eluted using two volumes of 25 µl of Qiagen EB buffer.

### Yeast transformation

The plasmid DNA, prepared as described above, was digested for one hour with 50 *Dpn*I (NEB) units to eliminate unreplicated plasmids, EtOH precipitated and resuspended in TE buffer. This DNA was used to transform yeast SMY537 strain (*MAT*a, *leu2-Δ1, trp1-Δ63, ura3–52, his3–200*, *bar1::HIS3*, can1::KanMX ^128^) by the lithium acetate method [3]. A fraction of each transformation mixture, 5%, was plated onto SC-Trp plates (synthetic complete, lacking tryptophan), and the remainder 95% onto 5-FOA-Trp (synthetic complete, containing 0.09% 5- FOA and lacking tryptophan) to score for expansions. Colonies on each plate were counted after three days of growth at 30°C. The frequency of expansions (confirmed by PCR as described below) was determined as the number of colonies obtained on SC-Trp-FOA divided by the total number of transformants on SC-Trp, with appropriate correction for dilution factors. A critical concern with this assay is to ensure that the expansion events occurred in the tissue culture cells and not during plasmid propagation in *E. coli*. We determine expansions-events baseline by transforming yeast cells directly with 100 ng of DNA plasmid preparation (without passing through mammalian cells). At least three independent transformants were tested to calculate the baseline and for each siRNA treatment.

### PCR analysis

To authenticate expansions and to determine their size, individual 5-FOA-resistant yeast colonies were disrupted with Lyticase as described in [4]. Subsequent PCR amplification by Phusion Polymerase (ThermoFisher) used UC1 (5’-GGTCCCAATTCTGCAGATATCCATCACAC-3’) and UC6 (5’-GCAAGGAATGGTGCATGCTCGAT-3’) primers flanking the repeat tract for 35 cycles of 20 s at 98°C and 2 min at 72°C with a final extension at 72°C for 4 min. The products were separated into 1.5% agarose gels. PCR product sizes were determined by comparison with 50 bp DNA ladder (New England Biolabs) using ImageLab^TM^ software (Bio-Rad). PCR confirmation of expansions is crucial since randomly occurring mutations within the *URA3* gene could result in a 5-FOA-resistant phenotype.

### Two-Dimensional (2D) agarose gel electrophoresis

Plasmid replication intermediates, extracted from human cells described above, were digested by *Xba*I*-Brs*GI-*Dpn*I restriction endonucleases (New England Biolabs), EtOH precipitated and resuspended in TE buffer. The first dimension was in a 0.4 % agarose gel in 1x TBE buffer (89 mM Tris-borate, 2 mM EDTA) at 1 V/cm at room temperature for 19 hours. The second dimension was in a 1% agarose gel in 1xTBE buffer and was run perpendicular to the first dimension. The dissolved agarose was poured around the excised agarose lane from the first dimension, and electrophoresis was at 5 V/cm in a 4°C cold chamber for nine hours. in the presence of 0.3 µg/ml ethidium bromide. Gels were washed for 15 min in 0.25 N HCl before an overnight transfer to a charged nylon membrane (Hybond-XL, GE Healthcare) in 0.4 N NaOH. The membrane was hybridized with a ^32^P-labeled radioactive probe corresponding to a 533 bp sequence comprising the *URA3* promoter and part of the CDS of the *URA3* gene. Membranes were washed sequentially twice with washing solution I (2xSSC, 1% SDS) at 65°C and twice with washing solution II (0.1×SSC, 0.1% SDS) at 42°C. Membranes were exposed on IR-sensitive screens for 1–5 days, and detection was performed on a Typhoon Imager (GE Healthcare). At least three independent transformants were tested for each siRNA knockdown.

### Statistical analysis

When indicated, statistical analysis was performed via Welch ANOVA test using GraphPad Prism version 8, GraphPad Software, San Diego, California USA.

## 6 SUPPLEMENTARY DATA

### SUPPLEMENTAL EXPERIMENTAL METHODS

#### Human gene knockdown by siRNAs

All ON-TARGETplus SMARTpool^®^ siRNAs (4 individual siRNAs combined) were purchased from Dharmacon: ATR (L-003201-00-0005), ATM (L-003202-00-0005), BRCA1 (L-003461-00-0005), BRCA2 (L-003462-00-0005), CLSPN (L-005288-00-0005), DDX11 (L-011843-00-0005), FANCJ (L-010587-00-0005) FEN1 (L-010344-00-0005), HLTF (L-006448-00-0005), POLD3(L-026692-01-0005), RAD51 (L-003530-00-0005), RAD52 (L-011760-00-0005), SHPRH (L-007167-00-0005), TIMELESS (L-019488-00-0005), ZRANB3 (L-010025-01-0005), Non-Targeting (D-001810-10-05), RECQ1 (L-013597-00-0005) WRN (L-010378-00-0005).

Antibodies to measure knockdown by Western blots were:

ATR (Cell Signaling Technology mAb #13934), ATM (Cell Signaling Technology mAb #2873), Brca1 (Invitrogen #MA123164), Brca2 (Millipore 05-666), Clspn (Abcam ab94945), Ddx11 (Abcam ab230017), FanJ (Abcam ab180853), Fen1 (Abcam ab17994), Hltf (Bethyl A300-230AM), PolD3 (POLD3 Bethyl Ab PolD3 A301-244A-T), Rad51 (Bioacademia 70-001), Rad52 (Acam ab 124971), Timeless (Abcam ab50943), Shprh (TrueMAB TA501443), Smarcal1 (Abcam6990), Zranb3 (Thermo 23111-1-AP), RECQ1 (Millipore ABC1428), WRN (Abcam ab17987).

## Acknowledgments

We thank Catherine Freudenreich, Mitch McVey, Karlene Cimprich, Alessandro Vindigni and members of the Mirkin laboratory for their important input on this project. We thank also Sanjukta Ghosh for her technical assistance. We are also grateful to Edvard Smith and Negin Mozafari, Karolinska Institutet, for valuable discussion of the experiments and results.

---

**Supplementary Figure 1.**
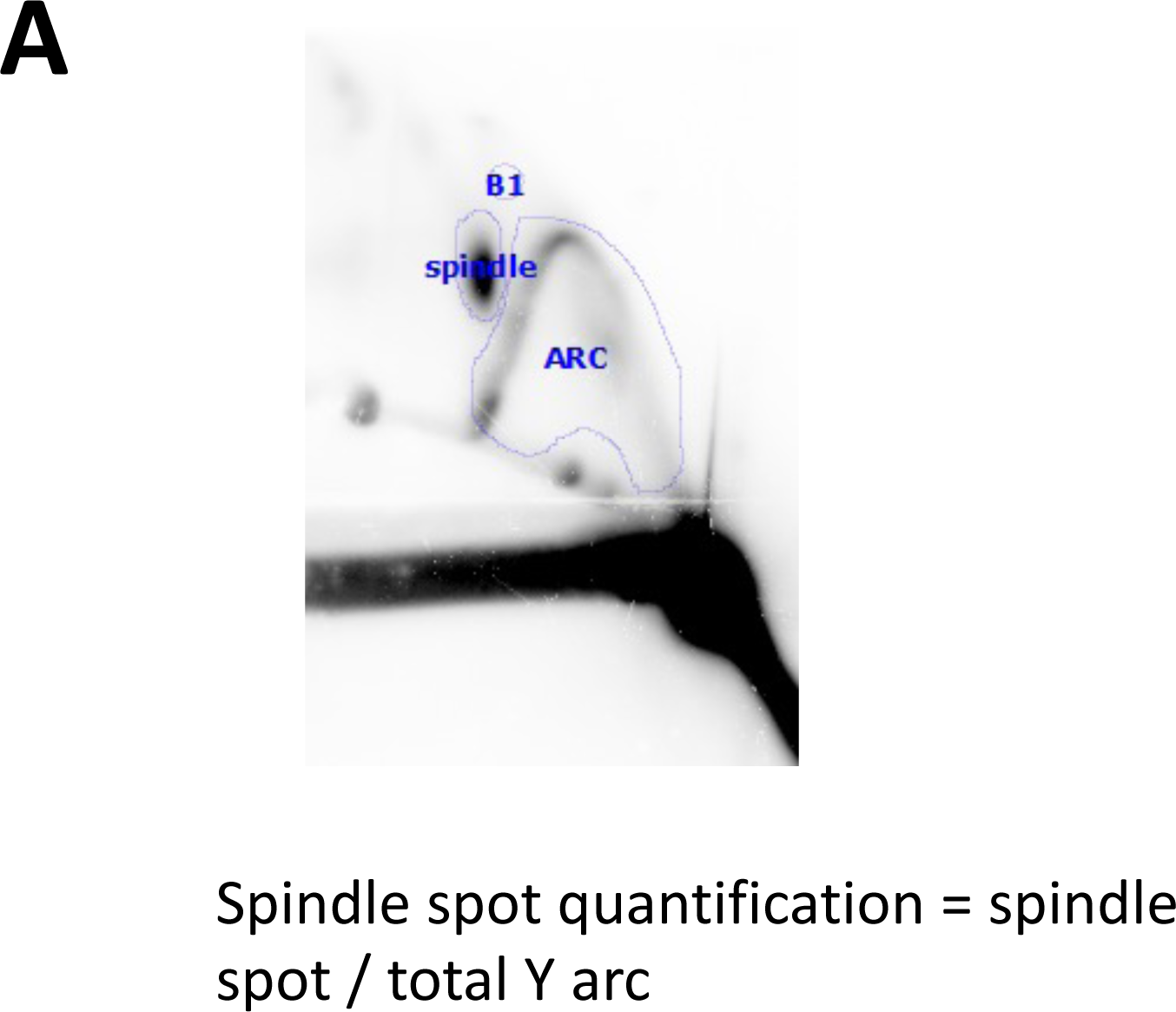
Quantification of the fraction of stalled forks and the angle of the arc deviation at the stall. (A) A representative 2D gel is shown with its densitometric profile corresponding to the Y-arc region where the GAA repeats are located; peaks on densitograms (red arrow) correspond to bulges on the Y-arcs. The ratio of radioactivity in the peak area to that corresponding area of a smooth replication arc reflects the fraction of stalled forks, and it is calculated as shown in the Figure. The densitometric profiles were obtained using the *plot-profile* tool available in the ImageJ (NIH) program (Schneider, C. A., Rasband, W. S., & Eliceiri, K. W. (2012). NIH Image to ImageJ: 25 years of image analysis. *Nature Methods*, *9*(7), 671–675. doi:10.1038/nmeth.2089). At least three experiments were analyzed for each siRNA treatment. **(B)** Spindle spot quantification using ImageLab®. Two circles were drawn; one represents the spindle spot intensity, and the other represents the background. The third volume represents the total arc of replication intermediates. Spindle spot quantification is made by dividing the intensity of the spindle spot compared to the full arc volume.

**Supplementary Figure 2.**
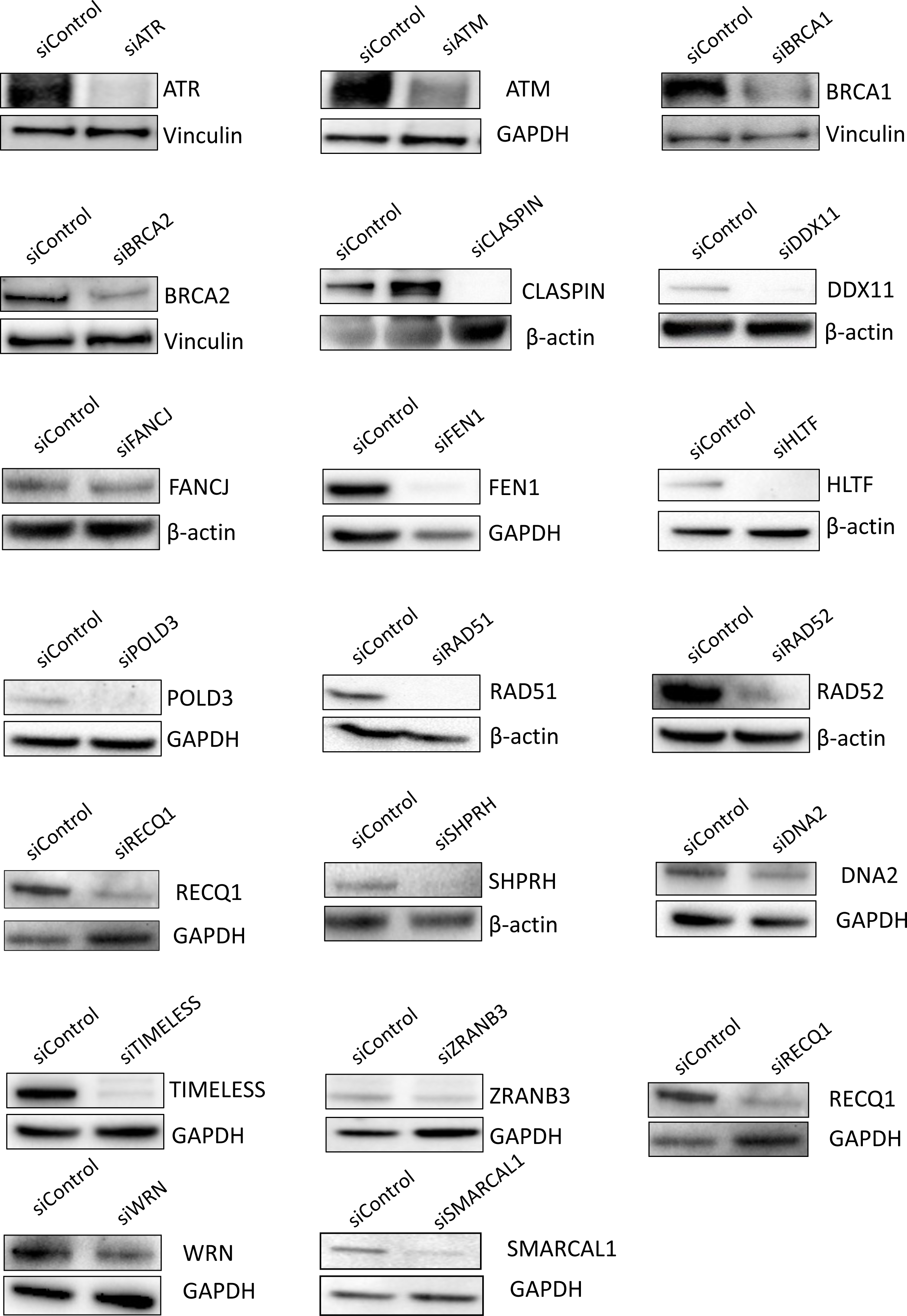
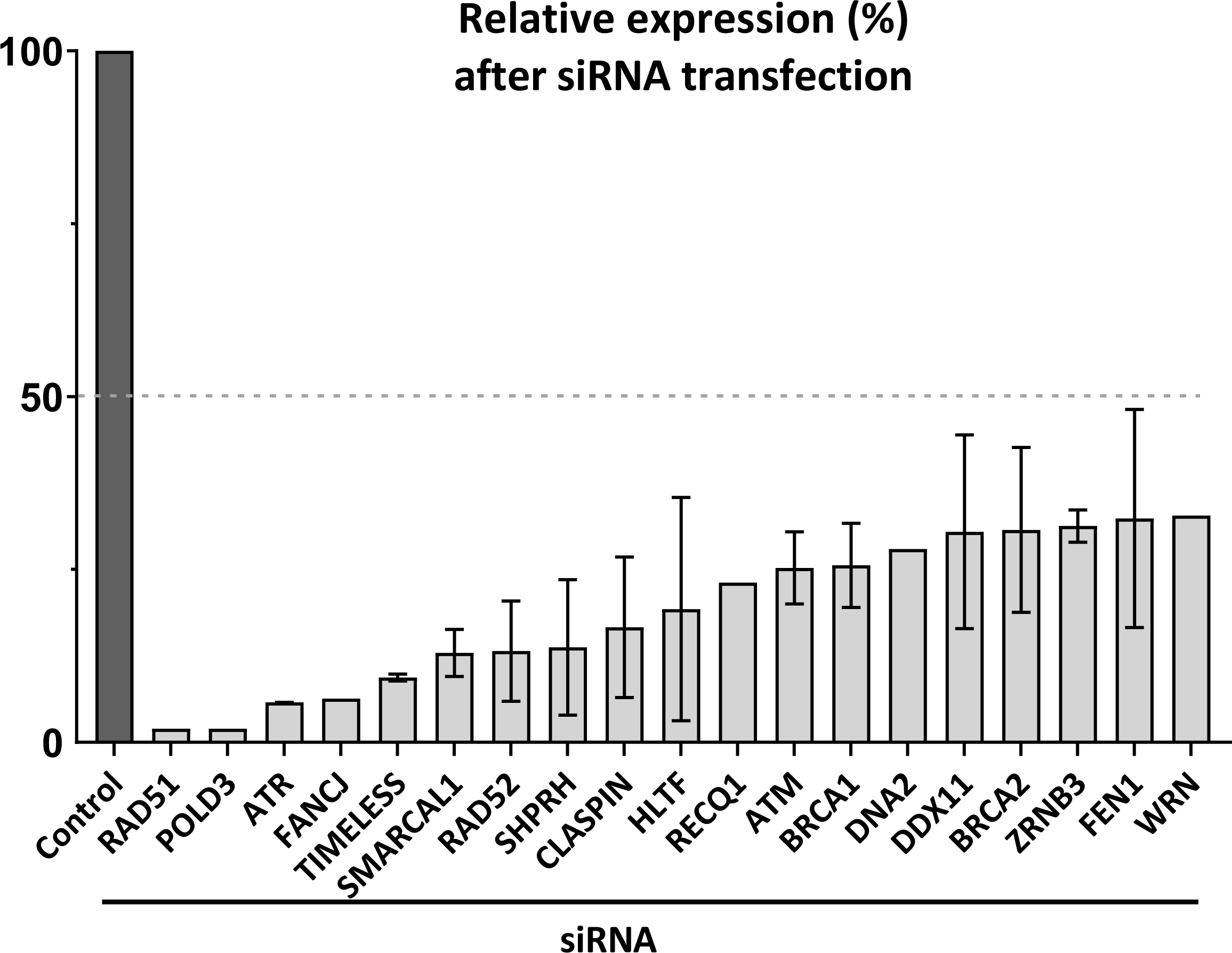
Validating the depletion of proteins via Western Blots. **(A)** Knockdown of human genes tested in this study by siRNAs. Illustrative Western Blot analyses with the indicated specific antibodies revealed marked reductions in the expression levels of the corresponding proteins. Protein levels for GAPDH, vinculin and β-actin are shown as an internal control. **(B)** Quantification of protein levels after using the corresponding pooled siRNAs.

**Supplementary Figure 3.**
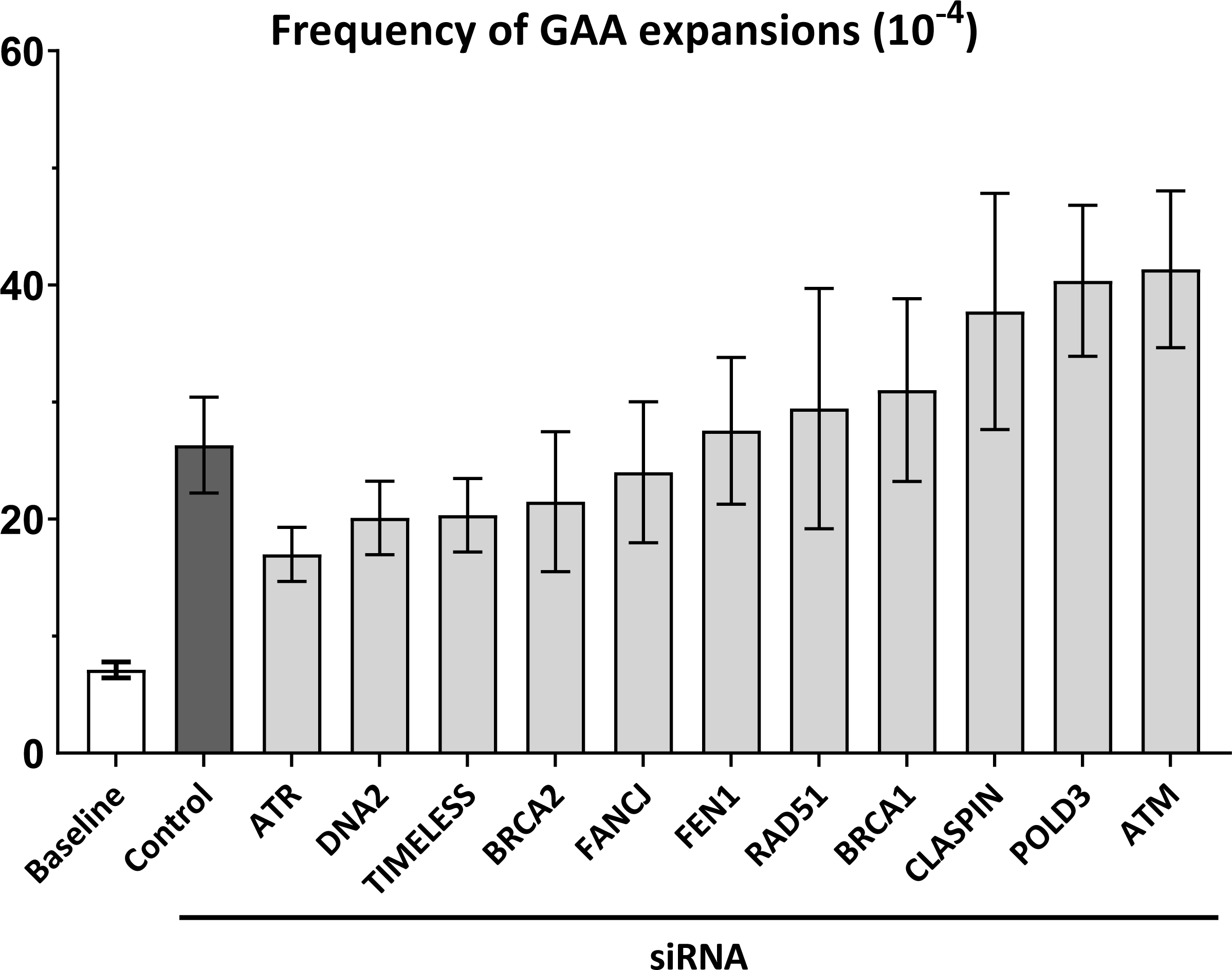
Frequencies of GAA repeat expansion after their replication in HEK-293T cells and upon candidate gene knockdown by siRNA with no statistical difference versus the control. The baseline expansion frequency upon yeast transformation by plasmid DNA isolated from *E. coli* is shown (white bar) for the comparison. Error bars indicate the standard error of the mean. Significance compared to the siControl frequency value was determined using a two-way Welch ANOVA test. See Table S1 for details.

**Supplementary Figure 4.**
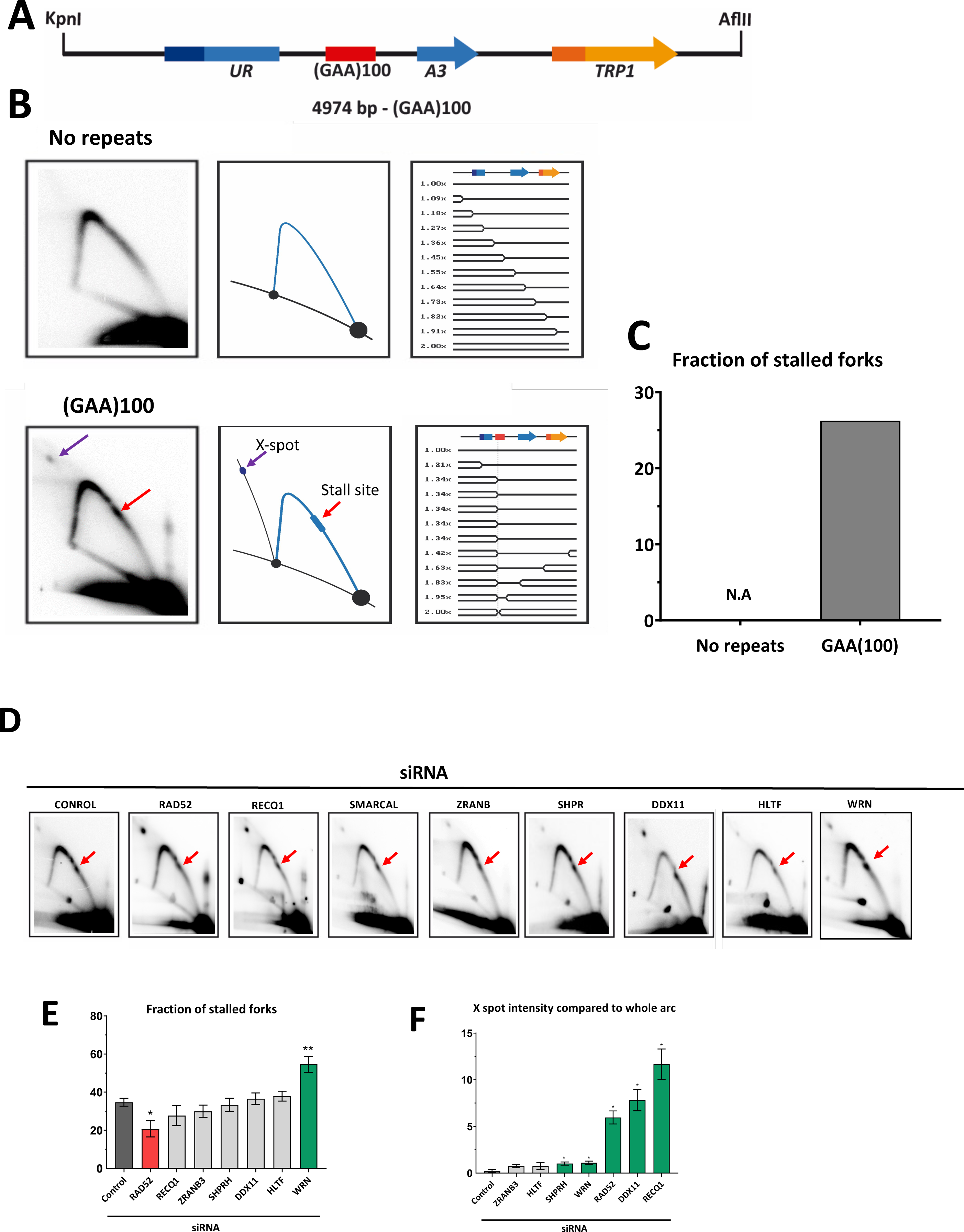

**Supplementary Figure 5.**
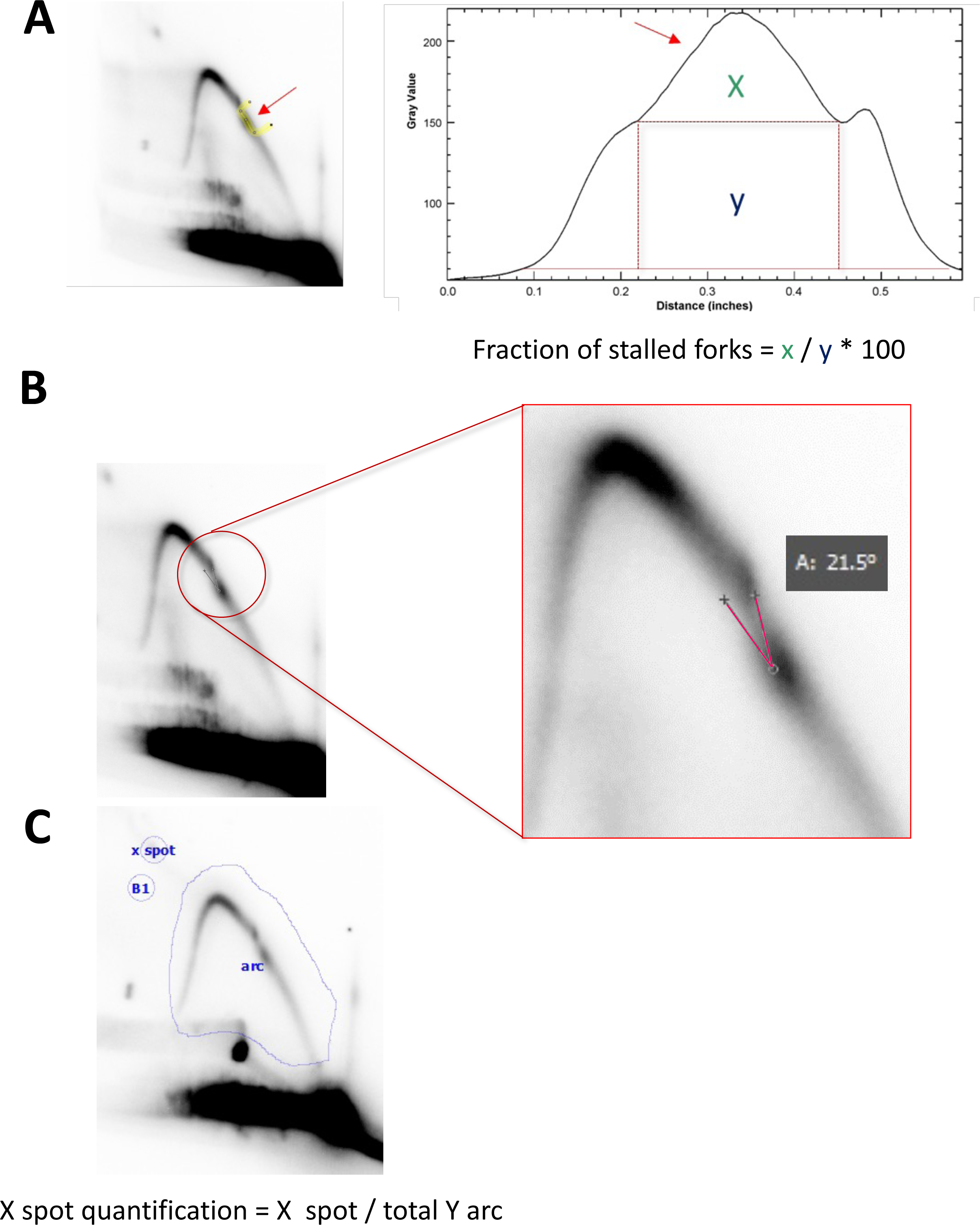

**Supplementary Table 1.**
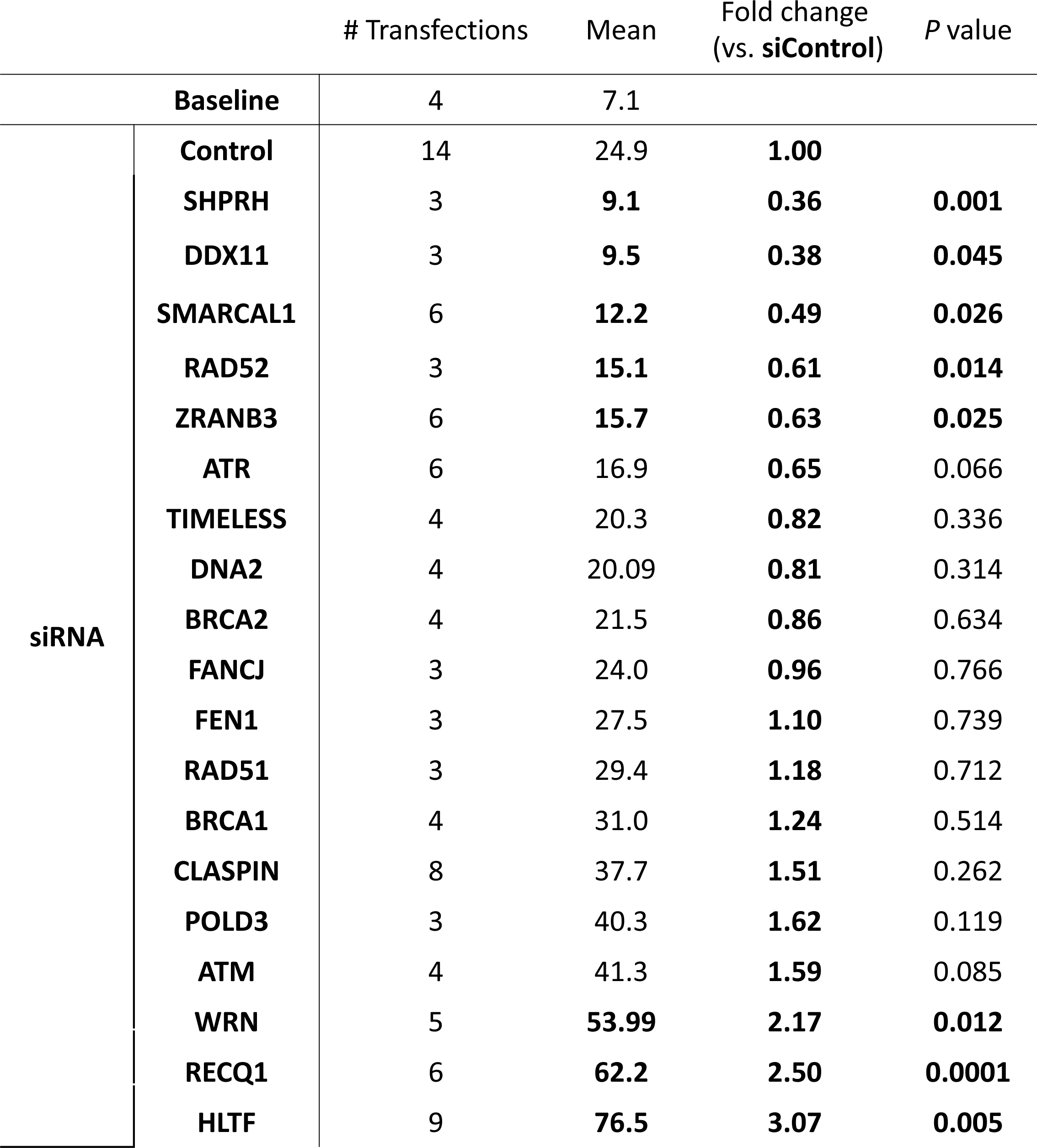
GAA repeat expansion frequencies.

**Supplementary Table 2.**
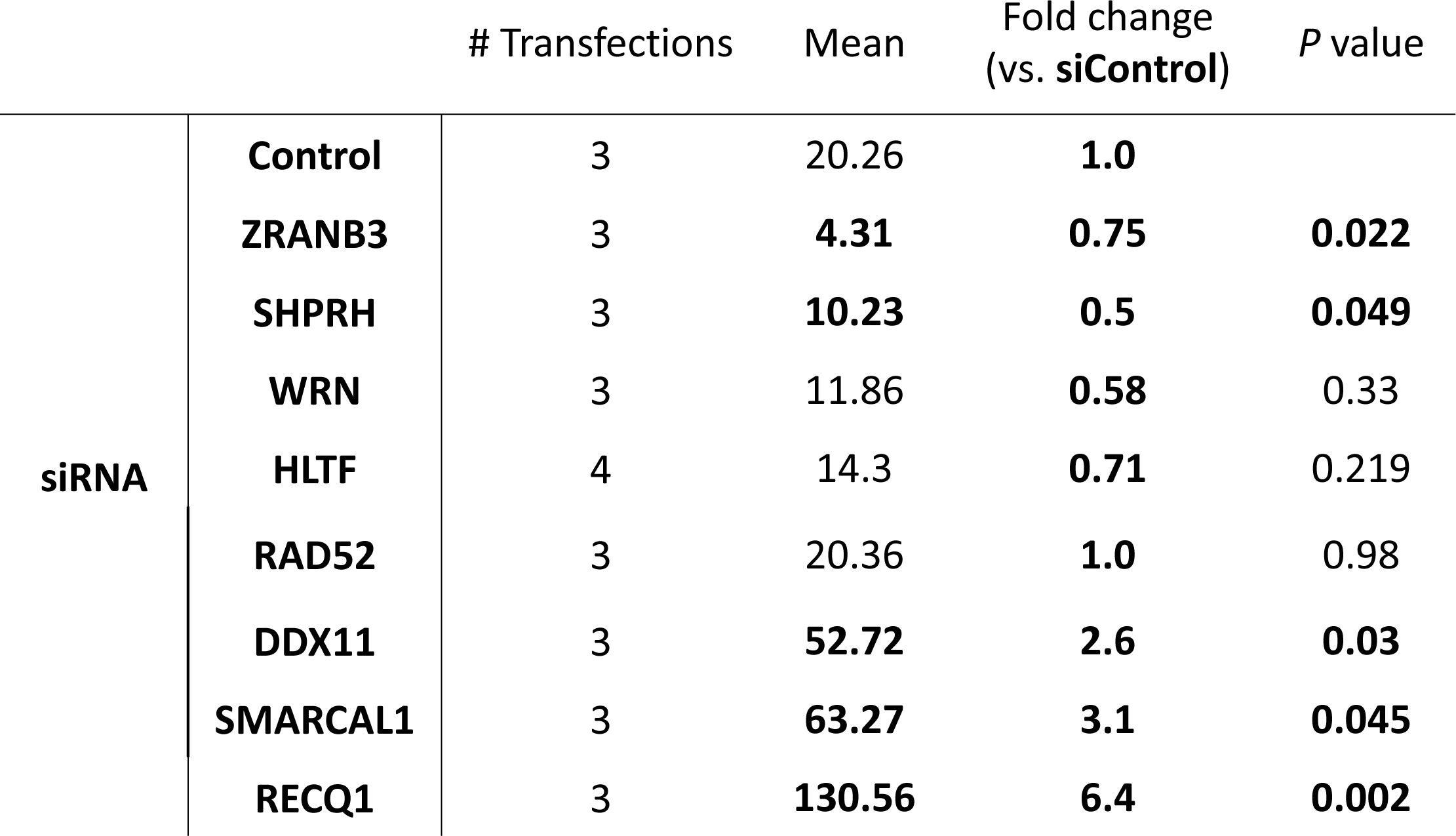
Quantification of spindle intermediates

|  |  | # Transfections | Mean | Fold change<br>(vs. siControl) | P value |
| --- | --- | --- | --- | --- | --- |
| siRNA | Control | 3 | 20.26 | 1.0 |  |
|  | ZRANB3 | 3 | 4.31 | 0.75 | 0.022 |
|  | SHPRH | 3 | 10.23 | 0.5 | 0.049 |
|  | WRN | 3 | 11.86 | 0.58 | 0.33 |
|  | HLTF | 4 | 14.3 | 0.71 | 0.219 |
|  | RAD52 | 3 | 20.36 | 1.0 | 0.98 |
|  | DDX11 | 3 | 52.72 | 2.6 | 0.03 |
|  | SMARCAL1 | 3 | 63.27 | 3.1 | 0.045 |
|  | RECQ1 | 3 | 130.56 | 6.4 | 0.002 |

**Supplementary Table 4.** Quantification of stalled forks

|  |  | # Transfections | Mean | Fold change<br>(vs. siControl) | P value |
| --- | --- | --- | --- | --- | --- |
| siRNA | Control | 3 | 34.7 | 1.00 |  |
|  | RAD52 | 3 | 20.7 | 0.60 | 0.048 |
|  | RECQ1 | 3 | 27.7 | 0.80 | 0.310 |
|  | ZRANB3 | 3 | 30.0 | 0.86 | 0.296 |
|  | RAD51 | 3 | 33.2 | 0.96 | 0.849 |
|  | SHPRH | 3 | 33.3 | 0.96 | 0.762 |
|  | DDX11 | 3 | 36.5 | 1.05 | 0.644 |
|  | HLTF | 3 | 37.9 | 1.09 | 0.392 |
|  | TIMELESS | 2 | 40.1 | 1.16 | 0.421 |
|  | FEN1 | 3 | 41.5 | 1.20 | 0.346 |
|  | CLASPIN | 6 | 44.3 | 1.28 | 0.153 |

**Supplementary Table 5.** Quantification of X intermediates

|  |  | # Transfections | Mean | Fold change<br>(vs. siControl) | P value |
| --- | --- | --- | --- | --- | --- |
| siRNA | Control | 6 | 0.23 | 1.0 |  |
|  | ZRANB3 | 3 | 0.75 | 3.3 | 0.06 |
|  | HLTF | 3 | 0.76 | 3.3 | 0.3 |
|  | SHPRH | 3 | 1.02 | 4.4 | 0.02 |
|  | RAD52 | 3 | 5.95 | 13.2 | 0.017 |
|  | DDX11 | 3 | 7.81 | 10.4 | 0.02 |
|  | RECQ1 | 3 | 11.66 | 15.3 | 0.02 |

